# Cell adhesion molecule IGPR-1 activates AMPK connecting cell adhesion to energy sensing and autophagy

**DOI:** 10.1101/2020.06.06.138214

**Authors:** Razie Amraei, Tooba Alwani, Rachel Xi-Yeen Ho, Zahra Aryan, Shawn Wang, Nader Rahimi

## Abstract

Immunoglobulin (Ig) and proline-rich receptor-1 (IGPR-1) is a cell adhesion molecule that regulates angiogenesis and endothelial barrier function. IGPR-1 is activated by shear stress and mediates endothelial cell’s response to shear stress. Autophagy plays critical roles in the maintenance of endothelial cells in response to cellular stress caused by blood flow. However, whether IGPR-1 is activated in response to, and mediates autophagy remains unknown. In this study, we demonstrate that IGPR-1 is activated by autophagy inducing stimuli, such as amino acid starvation, nutrient deprivation, rapamycin and lipopolysaccharide (LPS). We have identified IκB kinaseβ (IKKβ) as a key serine/threonine kinase activated by autophagy stimuli and mediates phosphorylation of IGPR-1 at Ser220. Activation of IGPR-1, in turn, stimulates phosphorylation of AMP-activated protein kinase (AMPK), which leads to phosphorylation of key pro-autophagy proteins, ULK1 and Beclin-1 (BECN1), increased LC3-II levels and accumulation of LC3 punctum. This study demonstrates that IGPR-1 is activated by and regulates autophagy, connecting cell adhesion to autophagy, a finding that has important significance for autophagy-driven pathologies such cardiovascular diseases and cancer.

## Introduction

Autophagy (also called macroautophagy), the lysosomal degradation of cytoplasmic organelles or cytosolic components, is an evolutionarily conserved cytoprotective mechanism that is induced in response to cellular stress, such as nutrient withdrawal, loss of cell adhesion, and flow shear stress, or by therapeutic genotoxic agents and others (1–4).

Upon induction of autophagy, unc-51-like kinase 1 (ULK1 also known as ATG1) associates with autophagy-related protein 13 (ATG13), and focal adhesion kinase family interacting protein of 200kD (FIP200) to form the ULK1 complex. ULK1 interaction with ATG13 and FIP200 is critical for ULK1 kinase activity and stability (5). The ULK1 complex translocates to autophagy initiation sites and recruits the class III phosphatidylinositol 3-kinase, vacuolar protein sorting 34 (VPS34) complex consisting of BECLIN-1 (the mammalian orthologue of the yeast autophagy protein Apg6/Vps30(6) and multiple other ATGs leading to the phagophore formation (7). The serine/threonine protein kinase mTOR (mechanistic target of rapamycin) complex 1 (mTORC1) is a key regulator of autophagy in response to nutrient availability. In the presence of amino acids, mTORC1 is activated and suppresses autophagy through phosphorylation of ULK1 and ATG13. However, upon nutrient deprivation, mTORC1 activity is inhibited, leading to the activation of ULK1 that induces the autophagy program (8,9). Suppression of mTORC1 activity by AMP-activated protein kinase (AMPK) is central to the regulation of autophagy. AMPK inactivates mTORC1 through phosphorylation of RAPTOR, a key protein present within the mTORC1 complex and more importantly directly phosphorylates ULK1 at multiple serine residues and activates it (10,11).

Commonly known autophagy inducing conditions or agents such as nutrient withdrawal including amino acid and serum starvation, immunosuppressant rapamycin/Sirolimus, and lipopolysaccharide (LPS) all activate several key kinases such as the IκB kinase (IKK) complex (12). IKK complex is composed of at least three proteins, including two catalytic subunits (IKKα and IKKβ) and the scaffold protein NF-κB essential modulator (NEMO; also called IKKγ) (13). In addition to its pivotal role in mediating phosphorylation of IκB(13), IKKβ can regulate autophagy in IκB-independent manner (14,15) by mechanisms that are not fully understood.

Immunoglobulin (Ig) and proline-rich receptor-1 (IGPR-1) was identified as a novel cell adhesion molecule expressed in various human cell types including, endothelial and epithelial cells and mediates cell-cell adhesion(16). IGPR-1 regulates angiogenesis, endothelial barrier function(16,17), decreases sensitivity of tumor cells to genotoxic agent, doxorubicin, and supports tumor cell survival in response to anoikis(18). IGPR-1 is localized to adherens junctions and, is activated through trans-homophilic dimerization(17). Additionally, IGPR-1 responds to various cellular stresses, as its phosphorylation (*i.e.,* Ser220) is significantly increased by flow shear stress(19) and exposure to doxorubicin(18,19). Curiously, both shear stress (20) and doxorubicin (21,22) are well-known potent inducer of autophagy, raising a possibility for the involvement of IGPR-1 in autophagy. In this study, we demonstrate that upon the induction of autophagy, IGPR-1 is phosphorylated at Ser220 via a mechanism that involves activation of IKKβ. IKKβ-dependent phosphorylation of IGPR-1 stimulates phosphorylation of AMPK, leading to activation of BECN1 and ULK1, connecting cell adhesion and energy sensing to autophagy.

## Results

### IGPR-1 is activated by autophagy

Homophilic transdimerization of IGPR-1 regulates its phosphorylation at Ser220 (23). Additionally, genotoxic agents such as doxorubicin(18) and flow shear stress (19) also simulate phosphorylation of IGPR-1at Ser220. As both shear stress and genotoxic agents are well-known for their roles in autophagy, we asked whether IGPR-1 is activated in response to autophagy. We used human embryonic kidney epithelial-293 (HEK-293) cells ectopically expressing IGPR-1 as a model system to study the role of IGPR-1 in autophagy, as these cells do not express IGPR-1 endogenously at the detectable level (24). To this end, we first, tested whether amino acid starvation, the best known inducer of autophagy, can stimulate phosphorylation of IGPR-1. Cells were lysed and whole cell lysates was subjected to Western blot analysis followed by immunoblotting with phospho-Ser220 and total IGPR-1 antibodies. Phosphorylation of IGPR-1 at Ser220 was significantly increased by brief amino acid starvation of HEK-293 cells. Increased in phosphorylation of Ser220 peaked after one minute with amino acid starvation and remained highly phosphorylated until 15 minutes (**Figure 1A**).

**Figure 1.**
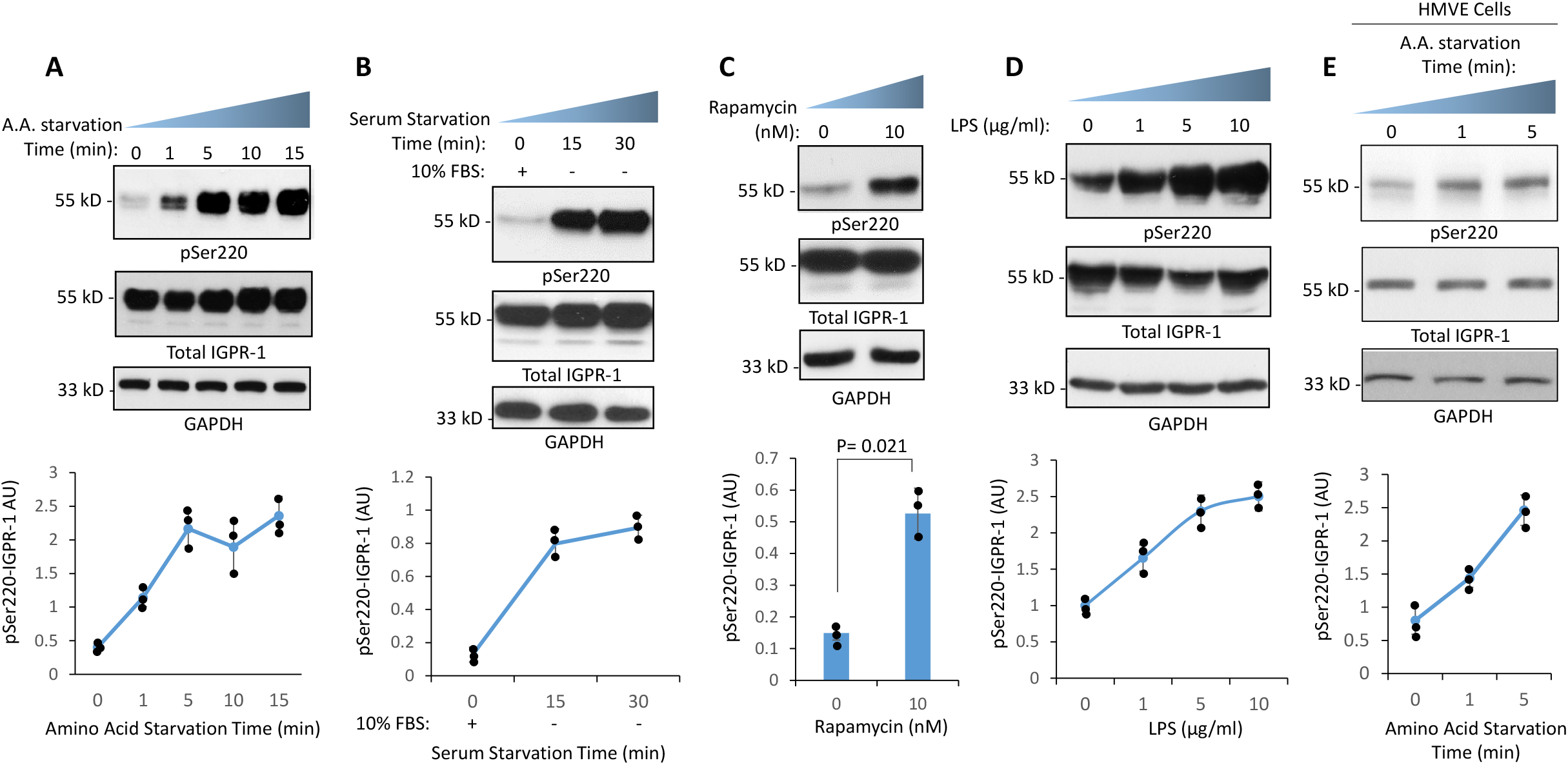
IGPR-1 is activated by autophagy stimuli. (**A**) IGPR-1/HEK-293 cells were kept in 10%FBS or in amino acid free media for indicated times. Whole cell lysates was prepared and subjected to Western blot analysis and blotted with anti-phosphoserine 220 IGPR-1 antibody (pSer220), anti-IGPR-1 antibody and anti-GAPDH antibody for protein loading control. Graph is a representative of three independent experiments. Individual data sets are shown (•). p<0.05. (**B**) IGPR-1/HEK-293 cells were seeded for 48hr in 10%FBS followed by serum starvation for 15 and 30 minutes by replacing the 10% DMEM media with serum-free DMEM media or cells left in 10% FBS DMEM media as a control (0). Cells were lysed and whole cell lysates blotted with the same antibodies as panel A. Graph is a representative of three independent experiments. Individual data sets are shown (•). p<0.05. (**C**) IGPR-1/HEK-293 cells were treated with vehicle (0) or with rapamycin (10nM) for one hour. Whole cell lysates prepared and subjected to Western blot analysis using the same antibodies as panel A. Graph is a representative of three independent experiments. Individual data sets are shown (•). P=0.021. (**D**) IGPR-1/HEK-293 cells were treated with different concentrations of LPS for one hour as indicated. Whole cell lysates was prepared and subjected to Western blot analysis and blotted with using the same antibodies as panel A. Graph is a representative of three independent experiments. Individual data sets are shown (•). p<0.05. (**E**) Human microvascular endothelial cells (HMVECs) were kept in 10%FBS or in amino acid free media for indicated times. Whole cell lysates was prepared and subjected to Western blot analysis and blotted with using the same antibodies as panel A. Individual data sets are shown (•). p<0.05.

Furthermore, in an additional set of experiments we subjected HEK-293 cells expressing IGPR-1 to various other autophagy inducing conditions or factors such as serum-starvation, rapamycin and LPS treatments and measured phosphorylation of Ser220. Both rapamycin and LPS are known to induce autophagy (12,25,26). Phosphorylation of IGPR-1 at Ser220 was significantly increased by brief serum-starvation of HEK-293 cells (**Figure 1B**). Furthermore, both rapamycin and LPS treatments of HEK-293 cells stimulated phosphorylation of IGPR-1 at Ser220 (**Figure 1C, D**). To demonstrate whether IGPR-1 is phosphorylated by autophagy in a biologically relevant human endothelial cells that IGPR-1 is expressed endogenously, we subjected primary human microvascular endothelial cells (HMVECs) to amino acid starvation. The result showed that IGPR-1 is phosphorylated at Ser220 in HMVECs (**Figure 1E**). Taken together, the data demonstrate that IGPR-1 is activated by autophagy in HEK-293 cells and human primary endothelial cells.

### IKKβ is activated by autophagy and phosphorylates IGPR-1

Activation of serine/threonine kinases represents a salient mechanistic feature of autophagy. Particularly, activation of IKKβ by nutrient deprivation (27), LPS(28,29) and rapamycin(30) is known to play a central role in autophagy. Therefore, we asked whether activation of IKKβ by serum-starvation, LPS or rapamycin can mediate phosphorylation of IGPR-1 at Ser220. To this end, we over-expressed wild type IKKβ or kinase inactive IKKβ (IKKβ-A44) in IGPR-1/HEK-293 cells. After 48 hours transfection, cells were either kept in 10% FBS or serum starved for 15, 30 or 60 minutes. Cells were lysed and whole cell lysates were immunoblotted for phospho-Ser220, total IGPR-1 and phospho-IKKβ. Expression of wild type IKKβ in IGPR-1/HEK-293 cells resulted in a robust phosphorylation of IGPR-1 in the presence of 10%FBS and this was further increased in response to serum starvation (**Figure 2A**). In contrast, over-expression of kinase inactive IKKβ-A44 in IGPR-1/HEK-293 cells markedly reduced phosphorylation of IGPR-1 at Ser220 (**Figure 2A**), indicating that IKKβ kinase activity is required for the serum-starvation dependent phosphorylation of IGPR-1 at Ser220. Similarly, LPS-induced phosphorylation of Ser220 on IGPR-1 was inhibited by a selective IKKβ inhibitor, IKK inhibitor III/BMS-345541(31) (**Figure 2B**). Furthermore, LPS stimulated phosphorylation of IKKβ and IKK inhibitor blocked its phosphorylation (**Figure 2B**). Conversely over-expression of a constitutively active IKKβ (IKKβ-S177E/S181E) in IGPR-1/HEK293 cells augmented LPS-induced phosphorylation of Ser220 (**Figure 2C**), indicating that IKKβ activity also is required for LPS-induced phosphorylation of IGPR-1 at Ser220.

**Figure 2:**
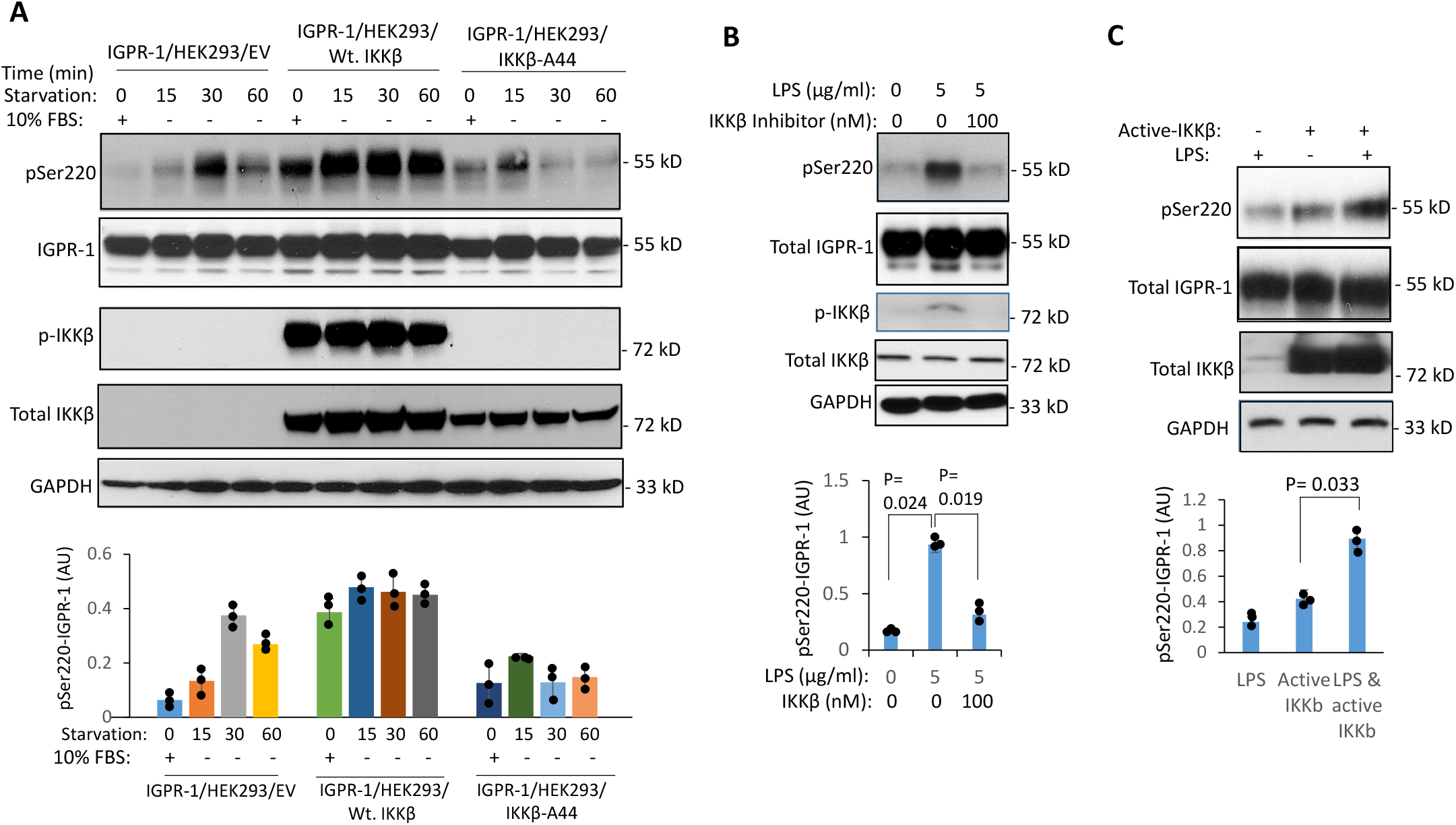
IKKβ mediates serum starvation-dependent and LPS-induced activation of IGPR-1. (**A**) IGPR-1/HEK293 cells were transfected with an empty vector (EV), wild-type IKKβ (Wt.IKKβ) or with kinase-dead IKKβ (IKKβ-A44). After 48 hours transfection, cells were either left in 10% FBS DMEM or serum starved for 15 or 30 minutes. Cells were lysed and whole cell lysates immunoblotted for pSer220, total IGPR-1, phospho-IKKβ, total IKKβ or blotted for GAPDH for protein loading control. Graph is a representative of three independent experiments. Individual data sets are shown (•). p<0.05. (**B**) IGPR-1/HEK-293 cells were treated with LPS (5μg/ml) alone or co-treated with LPS and IKKβ inhibitor, IKK inhibitor III/BMS-345541 (100ηM) for 60min. Cells were lysed and whole cell lysates was subjected to Western blot analysis and probed for pSer220, total IGPR-1, phospho-IKKβ, total IKKβ and GAPDH for protein loading control. Graph is a representative of three independent experiments. Individual data sets are shown (•). (**C**) IGPR-1/HEK-293 cells were transfected with an empty vector, EV (-) or constitutively active IKKβ (IKKβ-S177E/S181E). After 48 hours cells were either left untreated (-) or treated with LPS (5μg/ml) for 60min (+). Whole cell lysates was subjected to Western blot analysis and probed for pSer220, total IGPR-1, total IKKβ and GAPDH for protein loading control. Graph is a representative of three independent experiments. Individual data sets are shown (•). P=0.033.

### IGPR-1 is a substrate for IKKβ, and is phosphorylated by IKKβ *in vitro* and *in vivo*

Ser220 and the surrounding amino acids in IGPR-1 are strongly conserved both in human and non-human primates (**Figure 3A**), suggesting an evolutionary conserved mechanism for the phosphorylation of Ser220. IKKβ phosphorylates peptides with aromatic residues at the −2 position, hydrophobic residues at the +1 position, and acidic residues at the +3 position(32), suggesting that IKKβ is a likely candidate kinase involved in the phosphorylation of IGPR-1 at Ser220 (**Figure 3B**). Therefore, we asked whether IKKβ can phosphorylate IGPR-1 at Ser220 independent of the autophagy inducing factors like serum-starvation or LPS and rapamycin. We over-expressed wild-type IKKβ or kinase inactive IKKβ-A44 in IGPR-1/HEK-293 and 48 hours after transfection, cells were lysed and phosphorylation of IGPR-1 was determined by Western blot analysis. Over-expression of wild type IKKβ increased phosphorylation of IGPR-1, whereas kinase inactive IKKβ-A44 inhibited phosphorylation of IGPR-1 at Ser220 (**Figure 3C**). Similarly, IKKβ inhibitor inhibited both phosphorylation of IGPR-1 and IKKβ (**Figure 3D**).

**Figure 3:**
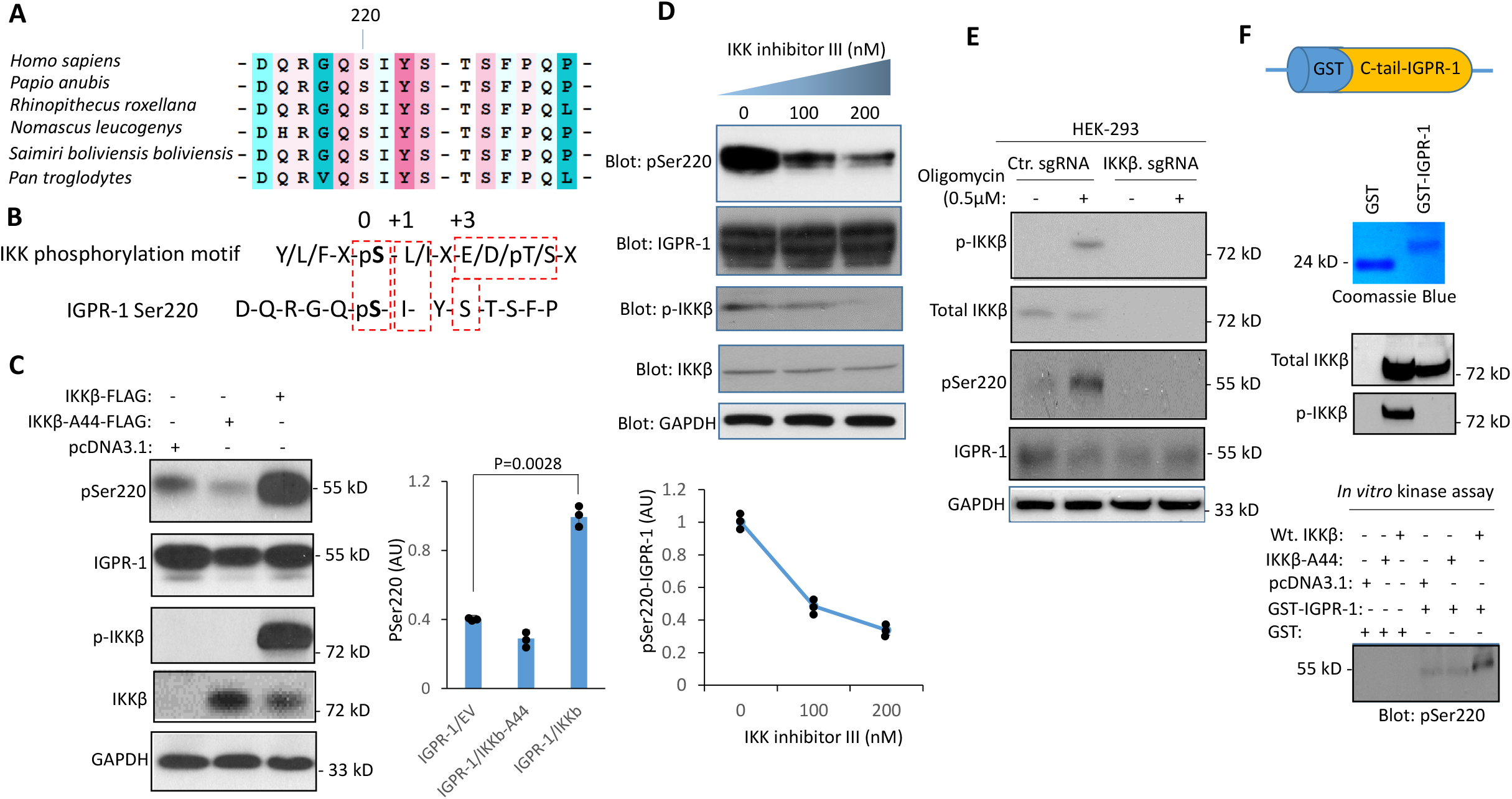
IGPR-1 is a substrate of and phosphorylated by IKKβ. (**A**) Ser220 and the surrounding amino acids on IGPR-1 in human and non-human primates is conserved. (**B**) IKKβ phosphorylation motif and IGPR-1 Ser220 phosphorylation site. (**C**) IGPR-1/HEK-293 cells were transfected with wild type IKKβ (Wt. IKKβ) or kinase inactive IKKβ (IKKβ-A44). After 48 hours cells were lysed and immunoblotted for pSer220, total IGPR-1, phospho-IKKβ, total IKKβ and GAPDH. Graph is a representative of three independent experiments. Individual data sets are shown (•). P=0.0028. (**D**) IGPR-1/HEK-293 cells were treated with IKK inhibitor III/BMS-345541 (100ηM) for 60min. Cells were lysed and immunoblotted for pSer220, total IGPR-1, phospho-IKKβ, and total IKKβ. Graph is a representative of three independent experiments. Individual data sets are shown (•). p<0.05. (**E**) Ctr.sgRNA/IGPR-1/HEK-293 cells and IKKβ-sgRNA/IGPR-1/HEK-293 cells were stimulated with IKKβ activator, Oligomycin (0.5μM), cells were lysed and subjected to Western blot analysis and immunoblotted for the indicated proteins. (**F**) In vitro kinase assay was performed by incubation of purified GST-cytoplasmic domain of IGPR-1 and purified wild type IKKβ or kinase inactive IKKβ-A44 in the presence of ATP. In vitro kinase reaction was stopped after 30min by heating the samples at 90°C for 5 min. Samples were subjected to Western blotting analysis and phosphorylation of Ser220 on GST-IGPR-1 was detected by immunoblotting with pSer220 antibody.

In an additional approach, we knocked out IKKβ via CRISPR-Cas9 system and examined the effect of loss of IKKβ in IGPR-1 phosphorylation. To this end, cells were either treated with a control vehicle or AMPK activator, Oligomycin and cells were lysed and phosphorylation of IGPR-1 at Ser220 was determined. Stimulation of IGPR-1/Ctr.sgRNA/HEK-293 cells with oligomycin stimulated AMPK activation and phosphorylation of IGPR-1 at Ser220 (**Figure 3E**). However, in IGPR-1/IKKβ.sgRNA/HEK-293 cells phosphorylation of Ser220 was not detected and treatment with oligomycin also did not stimulate phosphorylation of IGPR-1 at Ser220 (**Figure 3E**). IKKβ.sgRNA mediated knocked out IKKβ is shown (**Figure 3E**).

We next asked whether IKKβ can directly phosphorylates IGPR-1 at Ser220. To examine the direct involvement of IKKβ in catalyzing the phosphorylation of IGPR-1, we carried out an *in vitro* kinase assay using a purified recombinant GST-IGPR-1 protein that only encompasses the cytoplasmic domain of IGPR-1 and demonstrated that IKKβ phosphorylates IGPR-1 at Ser220 (**Figure 3F**). Taken together, our data identifies IGPR-1 as a novel substrate of IKKβ.

### IGPR-1 activates AMPK and stimulates phosphorylation of Beclin-1 and ULK1

We sought to determine whether IGPR-1 could activate AMPK, a widely considered master regulator of autophagy (33). HEK-293 cells expressing an empty vector (EV), IGPR-1 or A220-IGPR-1 were either maintained in 10%FBS or serum starved for 30 minutes or 12 hours. The Phospho-Thr172-AMPK immunoblot of whole cell lysates showed that expression of IGPR-1 in HEK-293 cells bypassed the requirement for serum starvation-dependent activation of AMPK as AMPK was strongly phosphorylated at Thr172 in IGPR-1/HEK-293 cells in 10%FBS compared to EV/HEK-293 cells (**Figure 4A**). Phosphorylation of AMPK was further augmented in IGPR-1/HEK-293 cells, particularly at 30min compared to control EV/HEK-293 cells (**Figure 4A**). Interestingly, A220-IGPR-1/HEK-293 cells showed also an increase in phosphorylation of AMPK, notwithstanding significantly less than the wild type IGPR-1, but more than the control EV/HEK-293 cells (**Figure 4A**). Phosphorylation of AMPKα at Thr172 in the activation loop is required for AMPK activation (34,35), indicating that expression of IGPR-1 in HEK-293 cells activated AMPK.

**Figure 4:**
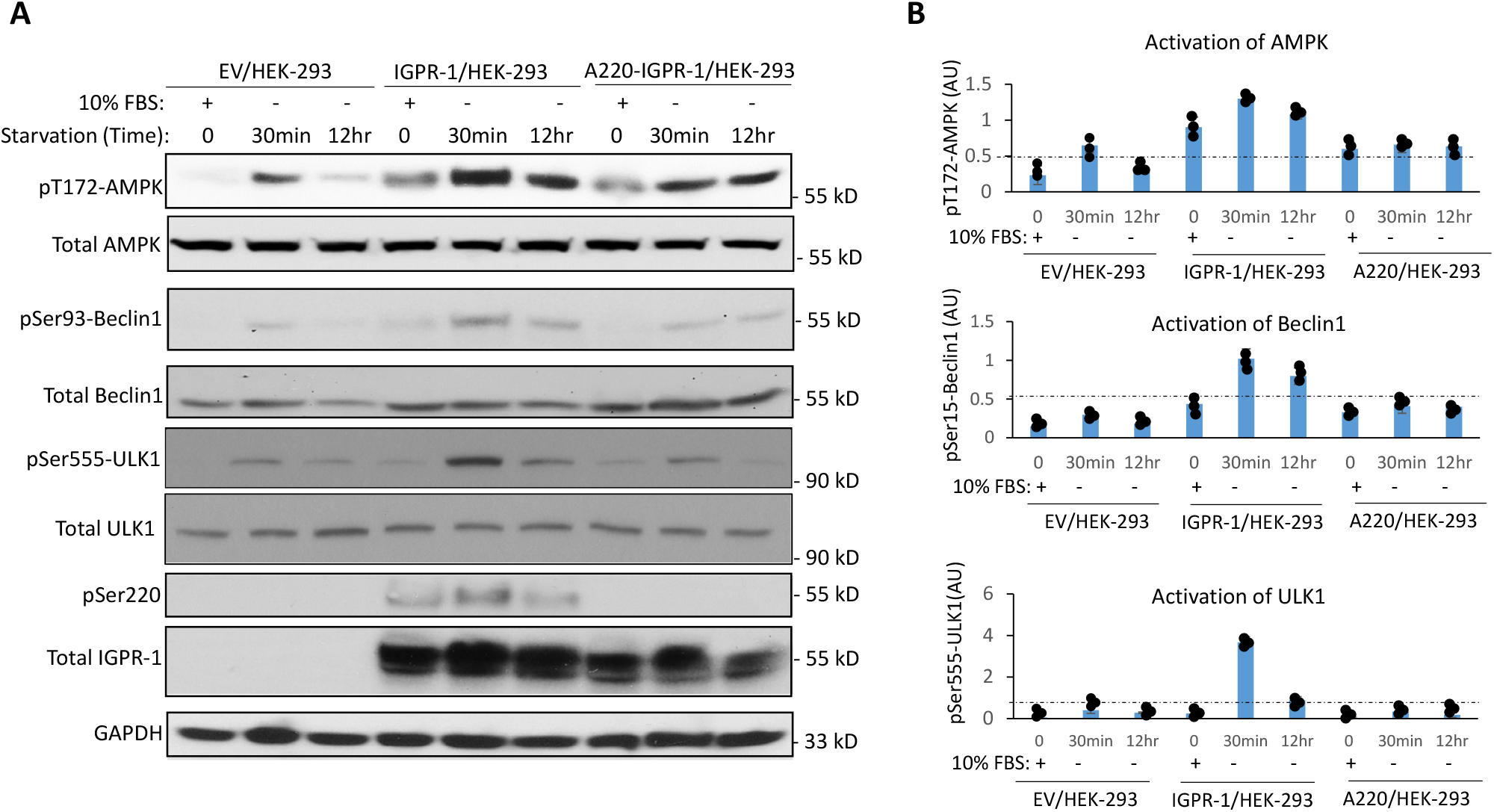
IGPR-1 stimulates phosphorylation of AMPK and mediates phosphorylation of ULK1. (**A**) HEK-293 cells expressing empty vector (EV), IGPR-1 or A220-IGPR-1 were either kept in 10%FBS DMEM or starved for 30 min or 12hours. Cells were lysed and immunoblotted for pT172-AMPK, pSer555-ULK1, pSer15-Beclin1, pSer220-IGPR-1, total IGPR-1 and GAPDH. (**B**) Graphs showing activation of AMPK, Beclin1 and ULK1 are representative of three independent experiments. Individual data sets are shown (•). p<0.05.

We next asked whether IGPR-1 induced AMPK activation also stimulates phosphorylation of Beclin-1 (BECN-1) and ULK1, which are the best-known substrates of AMPK and play central roles in autophagy(36). Similar to AMPK activation, phosphorylation of BECN1 at Ser93 was significantly increased in IGPR-1/HEK-293 cells compared to EV/HEK-293 or A220-IGPR-1 cells (**Figure 4A**), indicating that IGPR-1 induced activation of AMPK in HEK-293 cells also induced phosphorylation of BECN1 (**Figure 4A**) as previously reported (37). Furthermore, phosphorylation of ULK1 at Ser555 was also increased in IGPR-1/HEK-293 cells compared to control EV/HEK-293 cells. Interestingly, ULK1 phosphorylation was significantly reduced in A220-IGPR-1 cells, indicating that phosphorylation of Ser220, in part, is required for phosphorylation of ULK-1 (**Figure 4A**). The data indicate that phosphorylation of Ser220 on IGPR-1 plays an important role in the phosphorylation of AMPK, BECN-1 and ULK1.

### IGPR-1 induces autophagy in HEK-293 cells

We next examined the role of IGPR-1 in the autophagosome formation by measuring expression of LC3-phosphatidylethanolamine conjugate (LC3-II) and p62 endogenously expressed in HEK-293 cells, which are required for autophagosome development during autophagy(38), by Western blot analysis. HEK-293 cells expressing EV or IGPR-1 were either kept at 10%FBS or serum-starved for overnight. Cells were lysed and expression of LC3 and p62 levels was determined by immunoblotting with LC3 and p62 antibodies. While expression of LC3II was increased in response to serum-starvation in EV/HEK-293 cells, however expression of LC3II was significantly higher both in 10%FBS and serum-starved conditions in IGPR-1/HEK-293 cells (**Figure 5A**), indicating that IGPR-1 through increased in expression of LC3II regulates autophagosome formation. Additionally, as expected, p62 level was markedly decreased in response to serum-starvation (**Figure 5A**).

**Figure 5.**
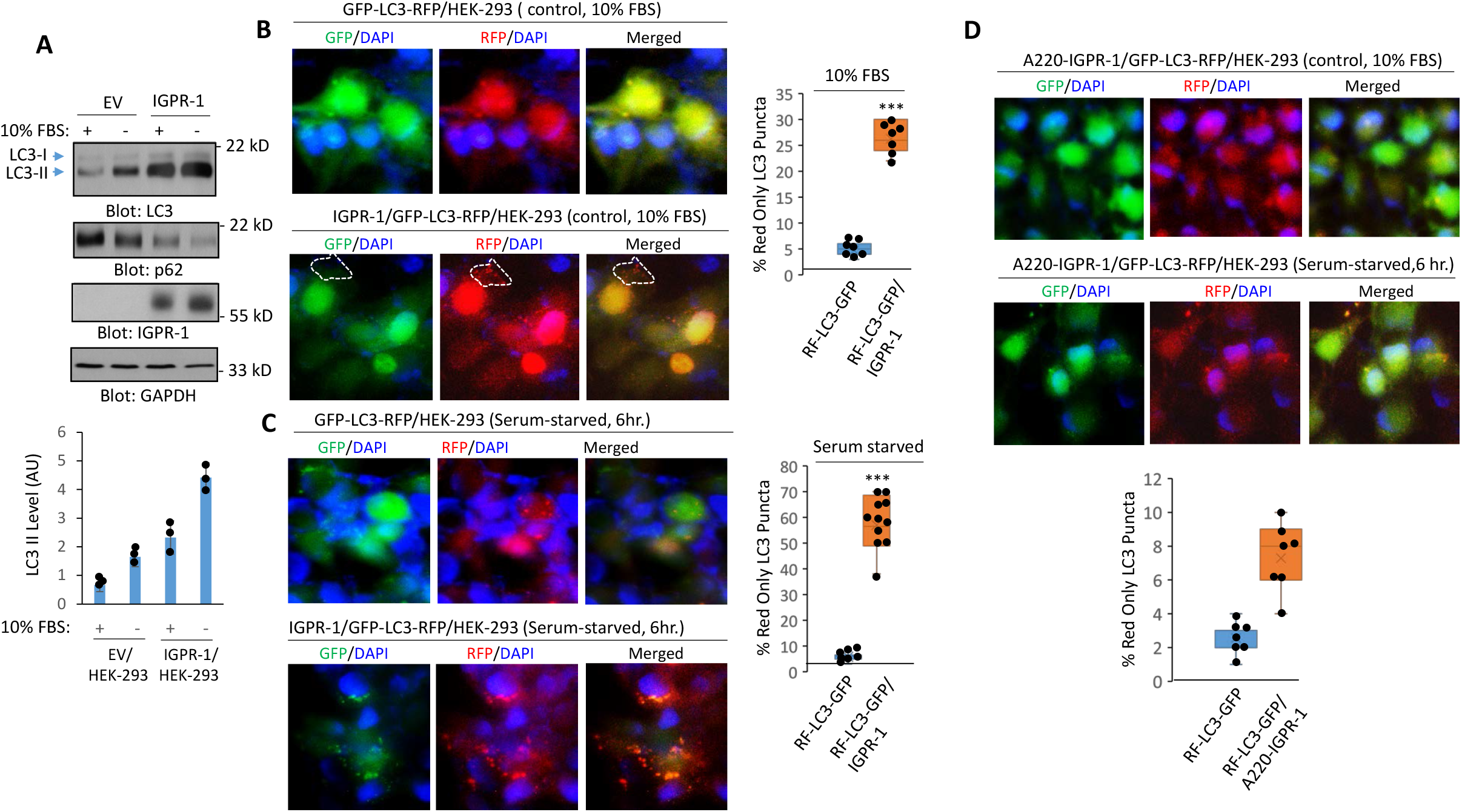
IGPR-1 mediates serum starvation-induced autophagy: (**A**) HEK-293 cells expressing empty vector (EV), IGPR-1 were either kept in 10%FBS or serum-starved for 12hours. Cells were lysed and immunoblotted for LC3 and p26, IGPR-1 and GAPDH. Graph is a presentative of three experiments. (**B**) HEK-293 cells expressing GFP-LC3-RFP/HEK-293 alone or co-expressing GFP-LC3-RFP and IGPR-1 either were kept in 10%FBS DMEM (**B**) or in serum-free DMEM for 6hours (**C**). Cells were fixed and stained with DAPI (nucleus) and viewed under a fluorescence microscope and pictures were taken. Image magnification (40X). (**C**) EV/HEK-293 and IGPR-1/HEK-293 cells were kept in 10% FBS or serum-starved for overnight, cells were lysed and whole cell lysates were blotted for LC3, p62, total IGPR-1, phospho-Ser220. Graph is a presentative of three independent experiments. Image J was used to quantify images (n=6 to 10 images for group). Individual data sets are shown (•). p<0.05. (**D**) HEK-293 cells co-expressing GFP-LC3-RFP and A220-IGPR-1 either were kept in 10%FBS DMEM or in serum-free DMEM for 6hours. Cells were fixed and stained with as panel B and viewed under a fluorescence microscope and representative pictures were taken. Image magnification (40X). Image J was used to quantify images (n=6 to 10 images for group). Individual data sets are shown (•). p<0.05.

To further elucidate the role of IGPR-1 in induction of autophagy, we established autophagic flux reporter cell lines by creating GFP-LC3 (microtubule-associated protein 1 light chain 3β)-RFP/HEK-293 and IGPR-1/GFP-LC3-RFP/HEK-293 cell lines via a retroviral expression system as previously reported(39). During autophagy, GFP-LC3-RFP labeled autophagosomes fuse with lysosomes. While the GFP signals are quenched due to the acidic environment (GFP is acid-sensitive) in the autolysosomes, the RFP signals remain stable as RFP is acid-stable and hence increased in number of RFP-LC3 (red only) puncta is considered a reflection of autophagic flux (40). In the presence of 10%FBS, only a few RFP-LC3 positive puncta were observed in GFP-LC3-RFP/HEK-293 cells (**Figure 5B**). However, we observed a significantly higher baseline of RFP-LC3 positive puncta, but not GFP-LC3 positive puncta, in IGPR-1/GFP-LC3-RFP/HEK-293 cells (**Figure 5C**). Moreover, when IGPR-1/GFP-LC3-RFP/HEK-293 cells were induced to undergo autophagy by serum starvation, they displayed a substantial increase in RFP-LC3 positive puncta (**Figure 5C**), suggesting that IGPR-1 regulates the formation of autophagosomes.

Next, we asked whether Ser220 mutant IGPR-1 (A220-IGPR-1) can induce RFP-LC3 puncta formation in HEK-293 cells. The result showed that the ability of A220-IGPR-1 to induce RFP-LC3 positive puncta was significantly reduced (**Figure 5D**). Altogether, the data demonstrate that IGPR-1 is activated by and regulates autophagy program.

## Discussion

Previous studies have shown that the activation of IKKβ regulates autophagy through mechanisms that involve expression of pro-autophagic genes via the NF-κB independent pathway and phosphorylation of the p85 subunit of PI3K, which leads to inhibition of mTOR (14,41). We revealed the existence of a previously unidentified pathway in autophagy which involves IKKβ-dependent activation of IGPR-1. We provide a mechanistic link between activation of IKKβ and phosphorylation of IGPR-1 at Ser220. IKKβ is activated by autophagy, leading to phosphorylation of IGPR-1 at Ser220 both in vivo and in vitro. Mechanistically, IKKβ mediated phosphorylation of IGPR-1 at Ser220 leads to activation of AMPK, which plays a central role in autophagy. Activation of IKKβ plays an essential role in autophagy as both the loss of function of IKKβ in mice and cell culture blocked autophagy (42). Likewise, IKKβ null cells are deficient in their ability to undergo autophagy in response to cellular starvation (15), which further underscores the critical role of IKKβ in autophagy.

AMPK is believed to exert its effect in autophagy by multiple mechanisms including, inactivating mTORC1 through phosphorylation of RAPTOR, a key protein present within the mTORC1, phosphorylating ULK1 at multiple serine residues including Ser555 that leads to its activation(10,11) and more importantly phosphorylating BECN1 at Ser93 and Ser96(37). IGPR-1 mediated activation of AMPK in HEK-293 cells increased phosphorylation of ULK1 at Ser555 and BECN1 at Ser93. Activation of AMPK and phosphorylation of BECN1 requires phosphorylation of IGPR-1 at Ser220. Activation of ULK1 and phosphorylation of BECN1 both play central roles in autophagy. ULK1 phosphorylation enables ULK1 to form a complex with ATG13 and focal adhesion kinase family interacting protein of 200 kd (FIP200) that leads to its translocation to autophagy initiation sites and subsequent recruitment of the class III phosphatidylinositol 3-kinase, vacuolar protein sorting 34 (VPS34) complex consisting of BECN1 and multiple other autophagy related proteins leading to the phagophore formation(6). Phosphorylation of BECN1 plays a key role in the initial steps in the assembly of autophagosomes from pre-autophagic structures is the recruitment and activation of VPS34 complex (36).

IGPR-1 is a cell adhesion molecule that mediates cell-cell adhesion and its activation regulates cell morphology and actin stress fiber alignment (24,43). The finding that IGPR-1 is activated by and regulates autophagy by stimulating activation of AMPK not only suggest a significant role for IGPR-1 in autophagy, but also links cell-cell adhesion to energy sensing and autophagy. Recent studies illuminated the key roles of autophagy in endothelial cells in response to various metabolic, blood flow-induced stresses and angiogenesis (44), the same cellular events also regulated by IGPR-1(24,43,45).

Additionally, cellular stress induced by flow shear stress (20,46) and exposure of cells to chemotherapeutic agent, doxorubicin (21,22) are linked to induction of autophagy, the conditions where IGPR-1 also is activated (18,45). Moreover, autophagy is associated with therapeutic resistance to chemotherapeutic agents (e.g., cisplatin, doxorubicin, temozolomide and etoposide), metabolic stresses, and small molecule inhibitors, suggesting a pro-tumor function for autophagy (47,48). Curiously, IGPR-1 is strongly phosphorylated by doxorubicin and regulates sensitivity of tumor cells to doxorubicin(18), indicating that IGPR-1 through induction of autophagy program could contribute to development of resistance in cancer cells.

Taken together, the data presented here suggest a significant role for IGPR-1 in autophagy and autophagy-associated diseases such as cancer and cardiovascular diseases. We propose IGPR-1 as a pro-autophagy cell adhesion molecule that upon activation stimulates AMPK activation, leading to phosphorylation of BECN1 and ULK1, key proteins involved in autophagy (**Figure 6**), linking cell adhesion to autophagy, a finding that has important significance for autophagy-driven pathologies such cardiovascular diseases and cancer.

**Figure 6.**
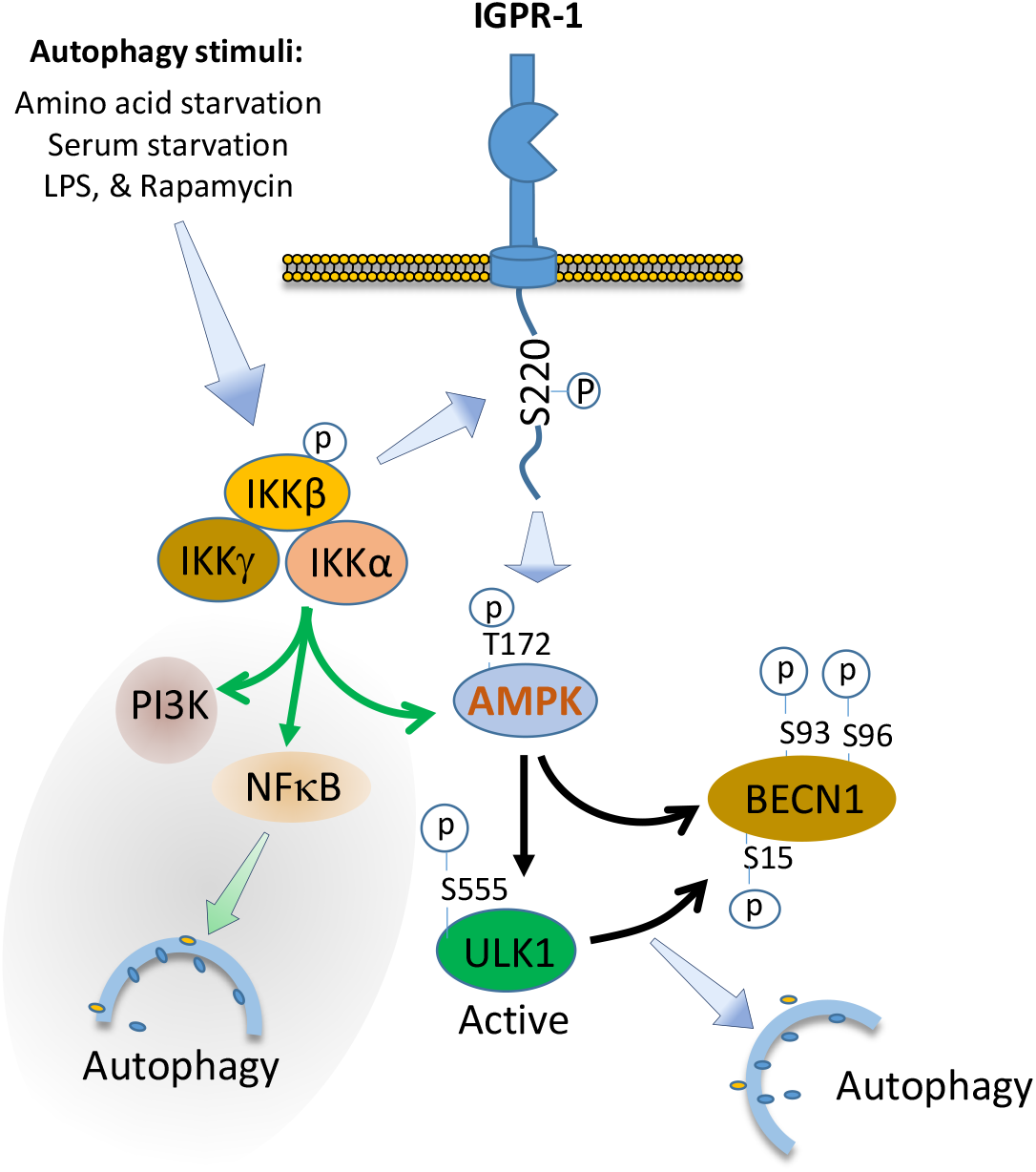
Proposed role of IGPR-1 in autophagy. Upon activation by autophagy stimuli, IGPR-1 is phosphorylated at Ser220 via IKKβ. Upon activation, IGPR-1 acts as a pro-autophagy signaling receptor leading to activation of AMPK. AMPK catalyze phosphorylation of BECN1 and ULK1, key proteins involved in autophagy. IGPR-1 is a dimeric protein and undergoes homophilic transdimerization in a cell density dependent manner (24) (not shown). Activation of IKKβ was previously thought to regulate autophagy by activation of AMPK, PI3K and induction of NFκB.

## Materials and Methods

### Antibodies, Plasmids, sgRNAs and chemicals

Anti–IGPR-1 and anti-phospho-Ser220 antibodies are homemade rabbit polyclonal antibodies previously described (24,43). Phospho-AMPK (T172), total AMPK, phospho-ULK1 (S555), total ULK1, phosph-Beclin-1 (S93), total Beclin-1, phospho-IKK (Ser176/180), total IKKβ, and LC3A/B, GAPDH antibodies all were purchased from Cell Signaling Technologies (Danvers, MA). The following plasmids were all purchased from Addgene (Watertown, MA); pcDNA3.FLAG-ULK1 (Cat#27636), pMRX-IP-GFP-LC3-RFP (Cat#84573), pcDNA-IKKβ-FLAG (Cat#23298), pcDNA-IKKβ-A44 (Cat#23299), and constitutive active IKKβ (S177E S181E, Cat#11105). IGPR-1 constructs including wild-type IGPR-1 and Ser220 mutant, A220-IGPR-1 constructs were cloned into retroviral vector, pQCXIP with C-terminal Myc tag as previously described (18,24,45). Retroviruses were produced in 293-GPG cells as described (49). IKK inhibitor III and rapamycin were purchased from Calbiochem and LPS was purchased from Sigma. Oligomycin was purchased from Cell Signaling Technologies (Danvers, MA). A set of three Human IKKβ sgRNAs (cat# GSGH-11938-16EG3551) were purchased from Dharmacon (Chicago, IL).

### Cell culture assays

HEK-293 cells expressing empty vector (EV), IGPR-1 or A220-IGPR-1 were maintained in DMEM supplemented with 10% fetal bovine serum and penicillin/streptomycin. To measure phosphorylation of IGPR-1 in response to serum-starvation, cells were plated in 60cm plates with 10%FBS DMEM for overnight with approximate 80-90% confluency. Cells were washed twice with PBS and cells were starved for 15 and 30 minutes or as described in the figure legends. Cells were lysed and whole cell lysates were mixed with sample buffer (5X) and boiled for 5minutes. Whole cell lysates were subjected to Western blot analysis and immunoblotted with antibody of the interest as described in the figure legends. In some experiments, cells were treated with a specific chemical inhibitor or transfected with a particular construct as indicated in the figure legends. Human microvascular endothelial cells were purchased from Cell Applications, INC., San Diego, CA and were grown in endothelial cell media.

### Recombinant GST fusion protein production

The generation of GST-fusion cytoplasmic domain of IGPR-1 cloned into pGEX-2T vector as previously described (43). The purified GST fusion IGPR-1 protein was subsequently used to measure the ability of IKKβ to phosphorylate IGPR-1 in an in vitro kinase.

### In vitro kinase assay

To detect phosphorylation of IGPR-1 at Ser220, the purified recombinant GST-IGPR-1 encompassing the cytoplasmic domain of IGPR-1 was mixed with wild type or kinase inactive IKKβ expressed in HEK-293 cells in 1× kinase buffer plus 0.2mM ATP, and incubated at 30°C for 15 min. The samples were mixed in 2X sample buffer and after boiling at 95°C for 5min were resolved on 12% SDS–PAGE followed by Western blot analysis using anti-phospho-Ser220 antibody.

### Western blotting analysis

The cells were prepared as described in the figures legends, lysed, and whole cell lysates were subjected to Western blot analysis. Normalized whole cell lysates were subjected to Western blotting analysis using IGPR-1 antibody, phospho-Ser220 antibodies or with appropriate antibody as indicated in the figure legends. Proteins were visualized using streptavidin–horseradish peroxidase–conjugated secondary antibody via chemiluminescence system. For each blot, films were exposed multiple times and films that showed within the linear range detection of protein bands were selected, scanned and subsequently used for quantification. Blots from at least three independent experiments were used for quantification purposes and representative data are shown. Image J software, an open source image processing program, used to quantify blots.

### Immunofluorescence microscopy

Cells expressing IGPR-1 or other constructs were seeded (1.5 × 10^6^ cells) onto coverslips and grown overnight in 60-mm plates to 90–100% confluence. The coverslips were mounted in Vectashield mounting medium with DAPI onto glass microscope slides. The slides were examined using a fluorescence microscope.

### Statistical analyses

Experimental data were subjected to Student t-test or One-way analysis of variance analysis where appropriate with representative of at least three independent experiments. p<0.05 was considered significant or as indicated in the figure legends.

## ACKNOWLEDGMENTS AND FUNDING

Funding: This work was supported in part through grants from the National institute of health NIH/NCI (R21CA191970, R21CA193958 and CTSI grant 1UL1TR001430 to N.R.).

## Author contributions

Razie Amraei, Tooba Alwani, Rachel Xi-Yeen Ho, Zahra Aryan and Shawn Wang and Nader Rahimi were involved in designing, performing and analyzing the experiments. Razei Amraie, Rachel Xi-Yeen Ho and Nader Rahimi were involved in writing of the manuscript.

## Data availability

Data and reagents are available from the corresponding author upon request

## References

1. Kroemer, G., Marino, G., and Levine, B. (2010) Autophagy and the integrated stress response. Molecular cell 40, 280–293

2. Vlahakis, A., and Debnath, J. (2017) The Interconnections between Autophagy and Integrin-Mediated Cell Adhesion. Journal of molecular biology 429, 515–530

3. Kenific, C. M., Wittmann, T., and Debnath, J. (2016) Autophagy in adhesion and migration. Journal of cell science 129, 3685–3693

4. Fung, C., Lock, R., Gao, S., Salas, E., and Debnath, J. (2008) Induction of autophagy during extracellular matrix detachment promotes cell survival. Mol Biol Cell 19, 797–806

5. Hurley, J. H., and Young, L. N. (2017) Mechanisms of Autophagy Initiation. Annu Rev Biochem 86, 225–244

6. Kametaka, S., Okano, T., Ohsumi, M., and Ohsumi, Y. (1998) Apg14p and Apg6/Vps30p form a protein complex essential for autophagy in the yeast, Saccharomyces cerevisiae. J Biol Chem 273, 22284–22291

7. Huang, W. P., and Klionsky, D. J. (2002) Autophagy in yeast: a review of the molecular machinery. Cell Struct Funct 27, 409–420

8. Jung, C. H., Jun, C. B., Ro, S. H., Kim, Y. M., Otto, N. M., Cao, J., Kundu, M., and Kim, D. H. (2009) ULK-Atg13-FIP200 complexes mediate mTOR signaling to the autophagy machinery. Mol Biol Cell 20, 1992–2003

9. Hosokawa, N., Hara, T., Kaizuka, T., Kishi, C., Takamura, A., Miura, Y., Iemura, S., Natsume, T., Takehana, K., Yamada, N., Guan, J. L., Oshiro, N., and Mizushima, N. (2009) Nutrient-dependent mTORC1 association with the ULK1-Atg13-FIP200 complex required for autophagy. Mol Biol Cell 20, 1981–1991

10. Kim, J., Kundu, M., Viollet, B., and Guan, K. L. (2011) AMPK and mTOR regulate autophagy through direct phosphorylation of Ulk1. Nat Cell Biol 13, 132–141

11. Egan, D. F., Shackelford, D. B., Mihaylova, M. M., Gelino, S., Kohnz, R. A., Mair, W., Vasquez, D. S., Joshi, A., Gwinn, D. M., Taylor, R., Asara, J. M., Fitzpatrick, J., Dillin, A., Viollet, B., Kundu, M., Hansen, M., and Shaw, R. J. (2011) Phosphorylation of ULK1 (hATG1) by AMP-activated protein kinase connects energy sensing to mitophagy. Science 331, 456–461

12. Kroemer, G., Marino, G., and Levine, B. (2010) Autophagy and the integrated stress response. Mol Cell 40, 280–293

13. Perkins, N. D. (2007) Integrating cell-signalling pathways with NF-kappaB and IKK function. Nature reviews Molecular cell biology 8, 49–62

14. Criollo, A., Senovilla, L., Authier, H., Maiuri, M. C., Morselli, E., Vitale, I., Kepp, O., Tasdemir, E., Galluzzi, L., Shen, S., Tailler, M., Delahaye, N., Tesniere, A., De Stefano, D., Younes, A. B., Harper, F., Pierron, G., Lavandero, S., Zitvogel, L., Israel, A., Baud, V., and Kroemer, G. (2010) The IKK complex contributes to the induction of autophagy. The EMBO journal 29, 619–631

15. Comb, W. C., Cogswell, P., Sitcheran, R., and Baldwin, A. S. (2011) IKK-dependent, NF-kappaB-independent control of autophagic gene expression. Oncogene 30, 1727–1732

16. Rahimi, N., Rezazadeh, K., Mahoney, J. E., Hartsough, E., and Meyer, R. D. (2012) Identification of IGPR-1 as a novel adhesion molecule involved in angiogenesis. Molecular biology of the cell 23, 1646–1656

17. Wang, Y. H. W., Meyer, R. D., Bondzie, P. A., Jiang, Y., Rahimi, I., Rezazadeh, K., Mehta, M., Laver, N. M. V., Costello, C. E., and Rahimi, N. (2016) IGPR-1 Is Required for Endothelial Cell-Cell Adhesion and Barrier Function. Journal of molecular biology 428, 5019–5033

18. Woolf, N., Pearson, B. E., Bondzie, P. A., Meyer, R. D., Lavaei, M., Belkina, A. C., Chitalia, V., and Rahimi, N. (2017) Targeting tumor multicellular aggregation through IGPR-1 inhibits colon cancer growth and improves chemotherapy. Oncogenesis 6, e378

19. Ho, R. X., Tahboub, R., Amraei, R., Meyer, R. D., Varongchayakul, N., Grinstaff, M. W., and Rahimi, N. (2019) The cell adhesion molecule IGPR-1 is activated by, and regulates responses of endothelial cells to shear stress. J Biol Chem

20. Liu, J., Bi, X., Chen, T., Zhang, Q., Wang, S. X., Chiu, J. J., Liu, G. S., Zhang, Y., Bu, P., and Jiang, F. (2015) Shear stress regulates endothelial cell autophagy via redox regulation and Sirt1 expression. Cell death & disease 6, e1827

21. Chen, H., Zhao, C., He, R., Zhou, M., Liu, Y., Guo, X., Wang, M., Zhu, F., Qin, R., and Li, X. (2019) Danthron suppresses autophagy and sensitizes pancreatic cancer cells to doxorubicin. Toxicol In Vitro 54, 345–353

22. Sui, X., Chen, R., Wang, Z., Huang, Z., Kong, N., Zhang, M., Han, W., Lou, F., Yang, J., Zhang, Q., Wang, X., He, C., and Pan, H. (2013) Autophagy and chemotherapy resistance: a promising therapeutic target for cancer treatment. Cell Death Dis 4, e838

23. Wang, Y. H., Meyer, R. D., Bondzie, P. A., Jiang, Y., Rahimi, I., Rezazadeh, K., Mehta, M., Laver, N. M., Costello, C. E., and Rahimi, N. (2016) IGPR-1 Is Required for Endothelial Cell-Cell Adhesion and Barrier Function. J Mol Biol 428, 5019–5033

24. Wang, Y. H. W., Meyer, R. D., Bondzie, P. A., Jiang, Y., Rahimi, I., Rezazadeh, K., Mehta, M., Laver, N. M. V., Costello, C. E., and Rahimi, N. (2016) IGPR-1 Is Required for Endothelial Cell-Cell Adhesion and Barrier Function. J Mol Biol 428, 5019–5033

25. Xu, Y., Jagannath, C., Liu, X.-D., Sharafkhaneh, A., Kolodziejska, K. E., and Eissa, N. T. (2007) Toll-like receptor 4 is a sensor for autophagy associated with innate immunity. Immunity 27, 135–144

26. Dunlop, E. A., and Tee, A. R. (2014) mTOR and autophagy: a dynamic relationship governed by nutrients and energy. Seminars in cell & developmental biology 36, 121–129

27. Comb, W. C., Hutti, J. E., Cogswell, P., Cantley, L. C., and Baldwin, A. S. (2012) p85alpha SH2 domain phosphorylation by IKK promotes feedback inhibition of PI3K and Akt in response to cellular starvation. Molecular cell 45, 719–730

28. Yang, F., Tang, E., Guan, K., and Wang, C. Y. (2003) IKK beta plays an essential role in the phosphorylation of RelA/p65 on serine 536 induced by lipopolysaccharide. J Immunol 170, 5630–5635

29. Dauphinee, S. M., and Karsan, A. (2006) Lipopolysaccharide signaling in endothelial cells. Lab Invest 86, 9–22

30. Dan, H. C., Cooper, M. J., Cogswell, P. C., Duncan, J. A., Ting, J. P. Y., and Baldwin, A. S. (2008) Akt-dependent regulation of NF-{kappa}B is controlled by mTOR and Raptor in association with IKK. Genes & development 22, 1490–1500

31. Burke, J. R., Pattoli, M. A., Gregor, K. R., Brassil, P. J., MacMaster, J. F., McIntyre, K. W., Yang, X., Iotzova, V. S., Clarke, W., Strnad, J., Qiu, Y., and Zusi, F. C. (2003) BMS-345541 is a highly selective inhibitor of I kappa B kinase that binds at an allosteric site of the enzyme and blocks NF-kappa B-dependent transcription in mice. The Journal of biological chemistry 278, 1450–1456

32. Hutti, J. E., Turk, B. E., Asara, J. M., Ma, A., Cantley, L. C., and Abbott, D. W. (2007) IkappaB kinase beta phosphorylates the K63 deubiquitinase A20 to cause feedback inhibition of the NF-kappaB pathway. Mol Cell Biol 27, 7451–7461

33. Kim, J., Kundu, M., Viollet, B., and Guan, K.-L. (2011) AMPK and mTOR regulate autophagy through direct phosphorylation of Ulk1. Nature cell biology 13, 132–141

34. Lizcano, J. M., Goransson, O., Toth, R., Deak, M., Morrice, N. A., Boudeau, J., Hawley, S. A., Udd, L., Makela, T. P., Hardie, D. G., and Alessi, D. R. (2004) LKB1 is a master kinase that activates 13 kinases of the AMPK subfamily, including MARK/PAR-1. EMBO J 23, 833–843

35. Hawley, S. A., Davison, M., Woods, A., Davies, S. P., Beri, R. K., Carling, D., and Hardie, D. G. (1996) Characterization of the AMP-activated protein kinase kinase from rat liver and identification of threonine 172 as the major site at which it phosphorylates AMP-activated protein kinase. J Biol Chem 271, 27879–27887

36. Menon, M. B., and Dhamija, S. (2018) Beclin 1 Phosphorylation - at the Center of Autophagy Regulation. Front Cell Dev Biol 6, 137

37. Kim, J., Kim, Y. C., Fang, C., Russell, R. C., Kim, J. H., Fan, W., Liu, R., Zhong, Q., and Guan, K. L. (2013) Differential regulation of distinct Vps34 complexes by AMPK in nutrient stress and autophagy. Cell 152, 290–303

38. Schaaf, M. B., Keulers, T. G., Vooijs, M. A., and Rouschop, K. M. (2016) LC3/GABARAP family proteins: autophagy-(un)related functions. FASEB J 30, 3961–3978

39. Kaizuka, T., Morishita, H., Hama, Y., Tsukamoto, S., Matsui, T., Toyota, Y., Kodama, A., Ishihara, T., Mizushima, T., and Mizushima, N. (2016) An Autophagic Flux Probe that Releases an Internal Control. Mol Cell 64, 835–849

40. Ni, H.-M., Bockus, A., Wozniak, A. L., Jones, K., Weinman, S., Yin, X.-M., and Ding, W.-X. (2011) Dissecting the dynamic turnover of GFP-LC3 in the autolysosome. Autophagy 7, 188–204

41. Comb, W. C., Hutti, J. E., Cogswell, P., Cantley, L. C., and Baldwin, A. S. (2012) p85alpha SH2 domain phosphorylation by IKK promotes feedback inhibition of PI3K and Akt in response to cellular starvation. Mol Cell 45, 719–730

42. Criollo, A., Senovilla, L., Authier, H., Maiuri, M. C., Morselli, E., Vitale, I., Kepp, O., Tasdemir, E., Galluzzi, L., Shen, S., Tailler, M., Delahaye, N., Tesniere, A., De Stefano, D., Younes, A. B., Harper, F., Pierron, G., Lavandero, S., Zitvogel, L., Israel, A., Baud, V., and Kroemer, G. (2010) The IKK complex contributes to the induction of autophagy. EMBO J 29, 619–631

43. Rahimi, N., Rezazadeh, K., Mahoney, J. E., Hartsough, E., and Meyer, R. D. (2012) Identification of IGPR-1 as a novel adhesion molecule involved in angiogenesis. Mol Biol Cell 23, 1646–1656

44. Nussenzweig, S. C., Verma, S., and Finkel, T. (2015) The role of autophagy in vascular biology. Circ Res 116, 480–488

45. Ho, R. X., Tahboub, R., Amraei, R., Meyer, R. D., Varongchayakul, N., Grinstaff, M., and Rahimi, N. (2019) The cell adhesion molecule IGPR-1 is activated by and regulates responses of endothelial cells to shear stress. J Biol Chem 294, 13671–13680

46. Guo, F., Li, X., Peng, J., Tang, Y., Yang, Q., Liu, L., Wang, Z., Jiang, Z., Xiao, M., Ni, C., Chen, R., Wei, D., and Wang, G. X. (2014) Autophagy regulates vascular endothelial cell eNOS and ET-1 expression induced by laminar shear stress in an ex vivo perfused system. Ann Biomed Eng 42, 1978–1988

47. Amaravadi, R. K., and Thompson, C. B. (2007) The roles of therapy-induced autophagy and necrosis in cancer treatment. Clin Cancer Res 13, 7271–7279

48. Degtyarev, M., De Maziere, A., Orr, C., Lin, J., Lee, B. B., Tien, J. Y., Prior, W. W., van Dijk, S., Wu, H., Gray, D. C., Davis, D. P., Stern, H. M., Murray, L. J., Hoeflich, K. P., Klumperman, J., Friedman, L. S., and Lin, K. (2008) Akt inhibition promotes autophagy and sensitizes PTEN-null tumors to lysosomotropic agents. J Cell Biol 183, 101–116

49. Rahimi, N., Dayanir, V., and Lashkari, K. (2000) Receptor chimeras indicate that the vascular endothelial growth factor receptor-1 (VEGFR-1) modulates mitogenic activity of VEGFR-2 in endothelial cells. J Biol Chem 275, 16986–16992

